# Limited Phylogenetic Overlap Between Fluoroquinolone-Resistant Escherichia coli Isolated on Dairy Farms and those Causing Bacteriuria in Humans Living in the Same Geographical Region

**DOI:** 10.1101/2021.04.24.441264

**Authors:** Oliver Mounsey, Hannah Schubert, Jacqueline Findlay, Katy Morley, Emma F. Puddy, Virginia C. Gould, Paul North, Karen E. Bowker, O. Martin Williams, Philip B. Williams, David C. Barrett, Tristan A. Cogan, Katy M. Turner, Alasdair P. MacGowan, Kristen K. Reyher, Matthew B. Avison

## Abstract

**Background:** Our primary aim was to test whether cattle-associated fluoroquinolone-resistant (FQ-R) *Escherichia coli* found on dairy farms were a significant cause of bacteriuria in humans living in the same 50 × 50 km geographical region located in South West England. Another aim was to identify risk factors for the presence of FQ-R *E. coli* on dairy farms.

**Methods:** FQ-R *E. coli* were isolated during 2017-18 from 42 dairy farms and from community urine samples. Forty-two cattle and 489 human urinary isolates were subjected to WGS, allowing phylogenetic comparisons. Risk factors were identified using a Bayesian regularisation approach.

**Results:** Of 489 FQ-R human isolates, 255 were also 3^rd^ generation cephalosporin-resistant (3GC-R), with strong genetic linkage between *aac(6’)Ib-cr* and *bla*_CTX-M-15_. We identified possible farm-to-human sharing for pairs of ST744 and ST162 isolates, but core genome SNP distances (71 and 63, respectively) were smaller in pairs of ST744 and ST162 isolates from different farms (7 and 3 SNPs, respectively). Total farm fluoroquinolone use showed a positive association with the odds of isolating FQ-R *E. coli* while total dry cow therapy use showed a negative association.

**Conclusions:** This work suggests that FQ-R *E. coli* found on dairy farms have a limited impact on community bacteriuria within the local human population, however, this appears greater than observed for 3GC-R *E. coli* when studied in parallel. Reducing fluoroquinolone use may reduce the on-farm prevalence of FQ-R *E. coli*, and this reduction may be greater when dry cow therapy is targeted to the ecology of resistant *E. coli* on the farm.

## Introduction

Fluoroquinolones are an important class of antibacterial drugs, included in the World Health Organisation’s list of ‘Highest Priority Critically Important Antimicrobials’.^1^ They are extensively used to treat infections in humans, companion animals, and farmed animals. These important medicines work by disrupting the activity of type II topoisomerases, causing the release of DNA that has double-strand breaks, leading to cell death.^2^

In addition to safety concerns about fluoroquinolone use in humans,^3^ bacteria such as *Escherichia coli* are increasingly becoming fluoroquinolone-resistant (FQ-R).^4^ Here, FQ-R is mainly caused by mutations altering the quinolone resistance-determining regions (QRDRs) of the primary target, DNA gyrase subunit A (GyrA), and the secondary target, DNA topoisomerase IV subunits (ParC) and, to a lesser extent, ParE.^5, 6^ However, additional mechanisms can also contribute to FQ-R, including regulatory mutations leading to AcrAB-TolC efflux pump over-production and expression of various plasmid-mediated quinolone resistance (PMQR) genes. For example, the various *qnr* genes, *oqxAB* efflux pump genes, and *aac(6’)Ib-cr*, which encodes a mutated aminoglycoside-modifying enzyme.^5, 7^

Because most FQ-R in *E. coli* results from multiple mutations in target enzymes and because PMQRs cannot confer FQ-R alone, the spread of FQ-R *E. coli* typically involves vertical dissemination of resistant clones. Most commonly in humans, the widespread proliferation of an FQ-R subset of the ST131 complex, ^8^ which is also frequently associated with 3^rd^ generation cephalosporin resistance (3GC-R). ^9^ Additionally, the pandemic ST1193 group,^10, 11^ which is only occasionally 3GC-R. ^10, 12^

In secondary care, the fluoroquinolone ciprofloxacin is commonly used as an oral switch following intravenous therapy for serious infections caused by Gram negative bacteria.^13^ Accordingly, FQ-R is an important threat to human health, and particularly when found in urinary *E. coli* since a substantial proportion of sepsis has an urinary origin and *E. coli* are the leading cause of urosepsis.^14^ The use of fluoroquinolones in primary care in the UK has reduced in recent years, primarily due to changes in policies concerning prescribing for community urinary tract infections. For example, in the 1.5 million population centred on the city of Bristol, we recently demonstrated a 24% fall in dispensing of ciprofloxacin, by far the most used fluoroquinolone in primary care, between 2013 and 2016. We noted a commensurate reduction in FQ-R in community-origin urinary *E. coli* in the same region.^15^

Fluoroquinolones have also been extensively used for the treatment of companion^16^ and farmed^17^ animals. Usage has decreased in farmed animals in the UK over recent years^18^ and voluntary use restrictions were introduced in June 2018 that mean fluoroquinolones are only used as a last resort, backed up by susceptibility testing^19^. There is still the possibility, however, that FQ-R *E. coli* that have been selected on farms might colonise humans and ultimately cause disease. There is also strong evidence that antibacterial resistant (ABR) *E. coli* are shared between companion animals and humans.^20^ Hence, considering FQ-R in humans through the lens of the One Health research framework may help obtain a wider picture of selection and transmission. Our primary aim was to test the hypothesis that FQ-R *E. coli* obtained in 2017-18 from dairy farms located within a 50 × 50 km region of South West England were closely related to FQ-R human urinary *E. coli* collected in parallel from the same region, suggestive of transmission. Whilst our similarly motivated studies of 3GC-R *E. coli* showed no evidence of recent sharing of isolates between dairy farms and humans,^21, 22^ we considered that the clonal nature of FQ-R *E. coli* might give a different outcome.

## Materials and Methods

### Isolates

Collection of FQ-R *E. coli* from dairy farms in South West England has been described previously.^23^ Briefly, 4145 samples were collected from faecally contaminated sites around 53 dairy farms (49 farms yielded FQ-R *E. coli*). Samples were collected at monthly visits between January 2017 and December 2018. FQ-R isolates were selected by plating processed samples onto Tryptone Bile X-Glucuronide (TBX) agar (Sigma-Aldrich, Dorset, UK) containing 0.5 mg/L ciprofloxacin, the EUCAST breakpoint.^24^ One FQ-R *E. coli* colony was picked from each plate if growth occurred. FQ-R urinary *E. coli* isolates were obtained from routine urine microbiology at Severn Infection Partnership, Southmead Hospital. Urine samples were submitted between Sept 2017 and Dec 2018 from 146 GPs located throughout Bristol and including coverage in Gloucestershire, Somerset, and Wiltshire.^15^ Bacterial identification was carried out using BD CHROMagar Orientation Medium (BD, GmbH, Heidelberg, Germany). Antibiotic susceptibilities were defined by disc testing using ciprofloxacin and ceftriaxone as indicators of FQ-R and 3GC-R as interpreted according to EUCAST guidelines.^24^ FQ-R/3GC-R urinary isolates and FQ-R but 3GC-susceptbile (FQ-R/3GC-S) isolates were all subcultured onto TBX agar containing 0.5 mg/L ciprofloxacin to confirm FQ-R status.

### Ethics

Farmers gave fully informed consent to participate in this study. Ethical approval was obtained from the University of Bristol’s Faculty of Health Sciences Research Ethics Committee (ref 41562). Ethical approval was not needed for the retention of human urinary isolates since these were anonymous and isolated for routine diagnostics.

### PCR, WGS, and Phylogenetic Analysis

PMQR genes (*qnrA, qnrB, qnrC, qnrD, qnrS, aac(6’)-Ib-cr, oqxAB*, and *qepA*) were identified by multiplex PCR as previously described.^25^ WGS was performed by MicrobesNG (https://microbesng.uk/) as previously described.^9, 22^ Resistance genes and STs were assigned using ResFinder ^26^ and MLST 2.0 ^27^ on the Centre for Genomic Epidemiology (http://www.genomicepidemiology.org/) platform.

Phylogenetic analysis was carried out using the Bioconda environment ^28^ on the Cloud Infrastructure for Microbial Bioinformatics.^29^ The reference sequences were ST131 isolate EC958 (accession: HG941718), ST744 isolate EC590 (accession: NZ_CP016182), and ST162 isolate W2-5 (accession: NZ_CP032989). Sequence alignments were with Snippy (https://github.com/tseemann/snippy). Maximum likelihood phylogenetic trees were constructed using RAxML, utilising the GTRCAT model of rate heterogeneity and the software’s autoMR and rapid bootstrap.^30, 31^ Trees were illustrated using Microreact.^32^

### Risk Factor Analysis

The risk factors fell into four categories: farm management, antibacterial usage (ABU), sample characteristics and meteorological. These categories, and the methods of data collection for each, are described in our recent wider risk factor study and its supplementary information.^23^ All code can be found at https://github.com/HannahSchubert1/OH-STAR-modelling-code. A random intercept Bayesian model with farm as a random effect was fitted using R with the BRMS (https://cran.r-project.org/web/packages/brms/index.html) package. The presence of FQ-R *E. coli* within a sample was the dependent variable. The remaining variables were all fixed effects and were split into two groups: “main effects” which were the ABU variables (measured as milligram per population-corrected unit) hypothesised to affect FQ-R *E. coli* presence, namely, total fluoroquinolone usage, novobiocin usage, and total 3^rd^ and 4^th^ generation cephalosporin usage; and “regularised effects” which were all remaining variables.

For the main effects, uninformative priors (normal distribution with a mean 0, standard deviation 5) were used; for the regularised effects, a regularizing prior (horseshoe prior with a single degree of freedom) ^33^ was used. The mean intercept was also given an uninformative prior (normal distribution with mean 0, standard deviation 5) while the standard deviation of the random effects was given the default prior for BRMS (a half-Student-t distribution with 3 degrees of freedom and a scale factor of 10). Four chains were sampled with 2000 iterations. The target acceptance criterion was increased from the default 0.8 to 0.98 to decrease the chance of sampling divergencies.

## Results

### Molecular epidemiology of FQ-R human urinary E. coli

A total of 489 FQ-R human urinary *E. coli* isolates were collected; 255 were also 3GC-R, 234 were 3GC-S. According to PMQR gene multiplex PCR, nineteen isolates harboured *qnr* genes (14 *qnrS*, 5 *qnrB*) and two carried *oqxAB*. Notably, 135 (52.9%) of the 3GC-R isolates but only 26 (11.1%) of 3GC-S isolates carried *aac(6’)Ib-cr* (Chi square: 96.7, p<0.0001).

WGS was performed for 188 FQ-R urinary isolates; 90 were 3GC-R and 98 were 3CG-S. **Figure 1** shows a phylogenetic tree drawn based on core genome alignment for these isolates. As well as illustrating the dominance of ST131 (mainly 3GC-R, green markers) and ST1193 (mainly 3GC-S, orange markers) this tree also shows that clusters of isolates fell into ST744, ST162, ST69 and ST38 (**Figure 1**). A total of 22 different combinations of QRDR mutations were identified in the 188 sequenced isolates. However, a large majority (155/188) fell within three QRDR types, identified here a Types A, B and C (**Table 1**). Isolates within common FQ-R clonal groups ST131 and ST1193 were dominated by QRDR Types A and B, respectively, whilst QRDR Type C was found in isolates from 13 diverse STs.

**Table 1.** Common QRDR mutation patterns identified within FQ-R human urinary *E. coli*.

| QRDR<br>Type | Associated ST | Mutations relative to wildtype strain<br>(K12) | Number of<br>isolates<br>(of 188) |
| --- | --- | --- | --- |
| Type A | ST131 | *gyrA* S83L, *gyrA* D87N, *parC* S80I,<br>*parC* E84V, *parE* I529L | 90 |
| Type B | ST1193 | *gyrA* S83L, *gyrA* D87N, *parC* S80I,<br>*parE* L416F | 37 |
| Type<br>C | Thirteen STs | *gyrA* S83L, *gyrA* D87N, *parC* S80I | 28 |

**Figure 1.**
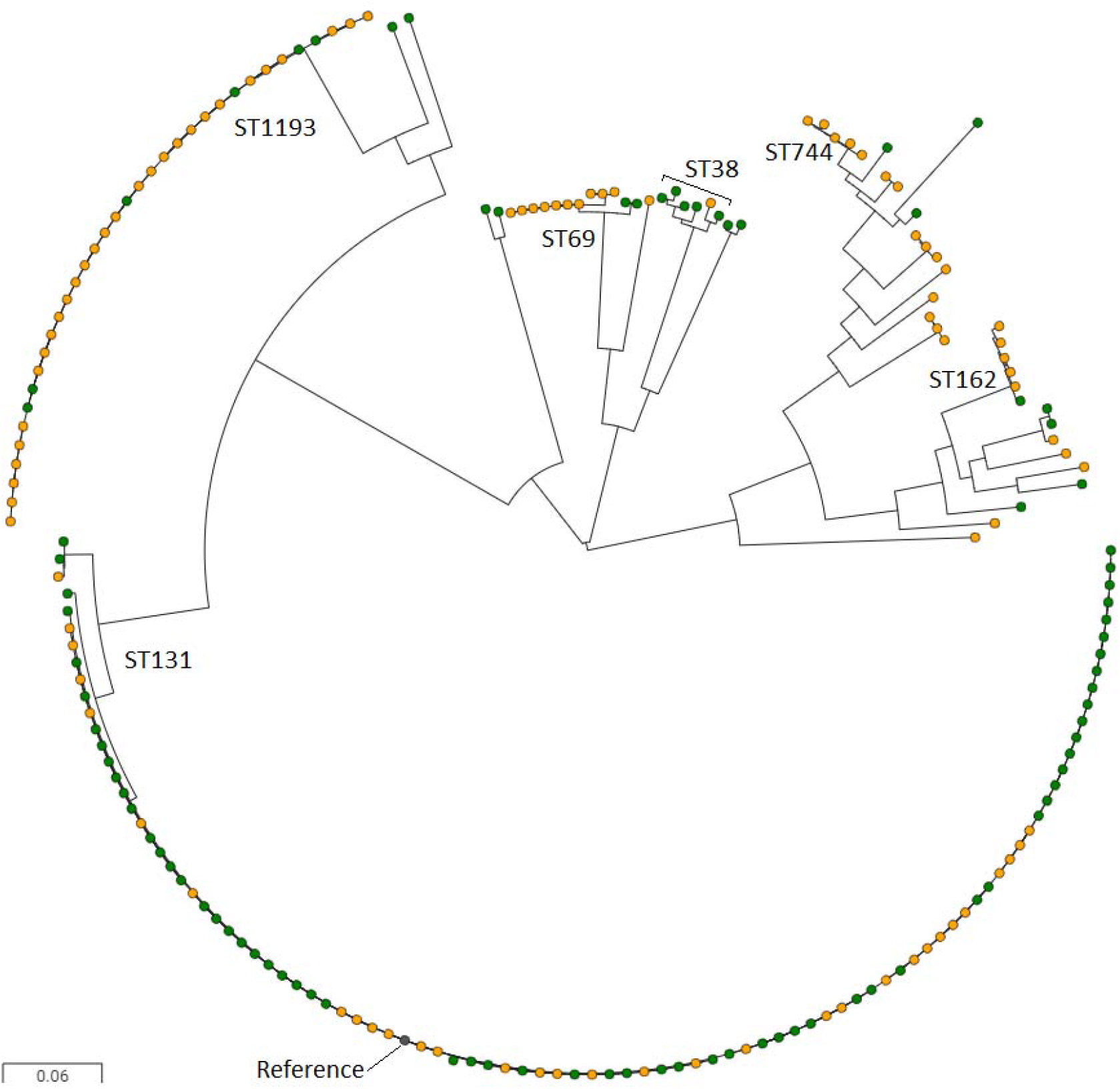
Maximum likelihood phylogenetic tree showing FQ-R urinary isolates. Green nodes were also 3GC-R, whilst yellow nodes were 3GC-S. Sequences were aligned against an ST131 reference (grey node). STs have been indicated where more than five isolates from that ST were present.

WGS also confirmed the strong bias for *aac(6’)Ib-cr* gene carriage, with 39 (43.3%) 3GC-R isolates being *aac(6’)Ib-cr*-positive, versus only four (4.1%) of the 3GC-S isolates (Chi square: 16.5, p<0.0001). We hypothesised that physical linkage of *aac(6’)Ib-cr* with an over-represented 3GC-R gene was driving this bias and **Table 2** reports that carriage of *aac(6’)Ib-cr* was strongly associated with carriage of *bla*_CTX-M-15_. We thus further hypothesised that this was due to physical linkage of *bla*_CTX-M-15_with the class 1 integron In*37*, carrying *aac(6’)Ib-cr*. Even given the difficulties of interpreting short-read WGS data, there was evidence in favour of this hypothesis. Of 11 isolates where the genomic locations of these genes could be definitively identified, in three, In*37* and *bla*_CTX-M-15_ were both embedded (at separate locations) in contigs surrounded by known chromosomal genes. In eight, In*37* and *bla*_CTX-M-15_ were immediately adjacent on the same contig, sometimes surrounded by known chromosomal genes, and sometimes not, suggesting a plasmid location.

**Table 2.** Chi square analyses examining the association of *aac(6’)Ib-cr* carriage with other different genotypic factors. All comparisons had degrees of freedom = 1.

| *aac(6')*/*lb-cr* carriage<br>versus: | Chi square | Association | p-value |
| --- | --- | --- | --- |
| ST131 | 6.2 | Positive | 0.013 |
| ST1193 | 4.3 | Negative | 0.038 |
| ST38 | 1.9 | Positive | 0.17 |
| *bla*<sub>CTX-M-15</sub> carriage | 47.8 | Positive | <0.0001 |
| *bla*<sub>CTX-M-27</sub> carriage | 5.0 | Negative | 0.025 |
| QRDR Type A | 8.7 | Positive | 0.003 |
| QRDR Type B | 3.5 | Negative | 0.060 |
| QRDR Type C | 3.0 | Negative | 0.083 |

3GC-R ST131 urinary *E. coli* isolates commonly carry *bla*_CTX-M-15_ in this study region.^9^ Therefore, it was not surprising to find a positive association between *aac(6’)Ib-cr* carriage and ST131 and between *aac(6’)Ib-cr* carriage and QRDR Type A (**Table 2**). However, of 32 FQ-R/3GC-S ST131 *E. coli*, only four carried *aac(6’)Ib-cr*, confirming *aac(6’)Ib-cr* was linked with *bla*_CTX-M-15_ and not with ST131.

In contrast, ST1193 isolates in this study were found mainly to be 3GC-S (n=28, versus n=6 for 3GC-R; **Figure 1**), and there was a significant negative association between *aac(6’)Ib-cr* carriage and the isolate being ST1193 (**Table 2**). There was also a negative association between carriage of *bla*_CTX-M-27_ and carriage of *aac(6’)Ib-cr* (**Table 2**). These associations reinforce the conclusion that In*37* and *bla*_CTX-M-15_ are closely genetically linked in urinary *E. coli* in our study region. No other association tested was found to be statistically significant.

### Molecular epidemiology of FQ-R E. coli from dairy farms and influence on human UTIs

We have recently reported a survey of resistant *E. coli* in faecal samples from 53 dairy farms in South West England.^23^ Of 4145 faecal samples collected from vicinities near animals, 263 were positive for FQ-R *E. coli*, representing 49 of 53 farms surveyed.^23^ Of 42 farms located within a 50 × 50 km region, which included the homes of the people providing the urinary samples discussed above, 245 FQ-R *E. coli* isolates from this earlier study ^23^ were taken forward for further analyses, representing one unique isolate from each positive sample. Multiplex PCRs showed that *aac(6’)Ib-cr* (found to be highly prevalent in the human urinary *E. coli* isolates) was entirely absent from the farm isolates and only nine isolates carried a PMQR gene (5 *qnrS*, 2 *qnrA* and 2 *qnrD)*.

WGS was performed for 42 FQ-R cattle-associated *E. coli* which were selected so that at least one isolate represented each farm that was positive for FQ-R *E. coli* within the 50 × 50 km region. Nine different STs were identified, with dominance of ST744 (21/42 isolates), ST162 (8/42 isolates), and ST10 (5/42 isolates). ST162 and ST10 have QRDR Type C, with three mutations as defined above (**Table 1**). ST744 also has these same three QRDR mutations, plus the addition of a distinctive *parC* mutation causing A56T. Because ST744 and ST162 were also common among FQ-R/3GC-S human isolates (**Figure 1**), we generated a phylogenetic tree to consider relationships between the cattle FQ-R and human FQ-R/3GC-S *E. coli* collected in parallel within our 50 × 50 km study area (**Figure 2**). This tree suggested overlapping ST744 and ST162 isolates found on different farms, but, importantly, also among human urinary isolates. Detailed trees generated using ST744 or ST162 reference genomes (**Figure 3**) confirmed this. The closest relationship between two human urinary isolates was 931 or 175 SNPs for ST744 and ST162, respectively, but the closest relationship between a human and a cattle isolate was 71 or 63 SNPs, respectively.

**Figure 2.**
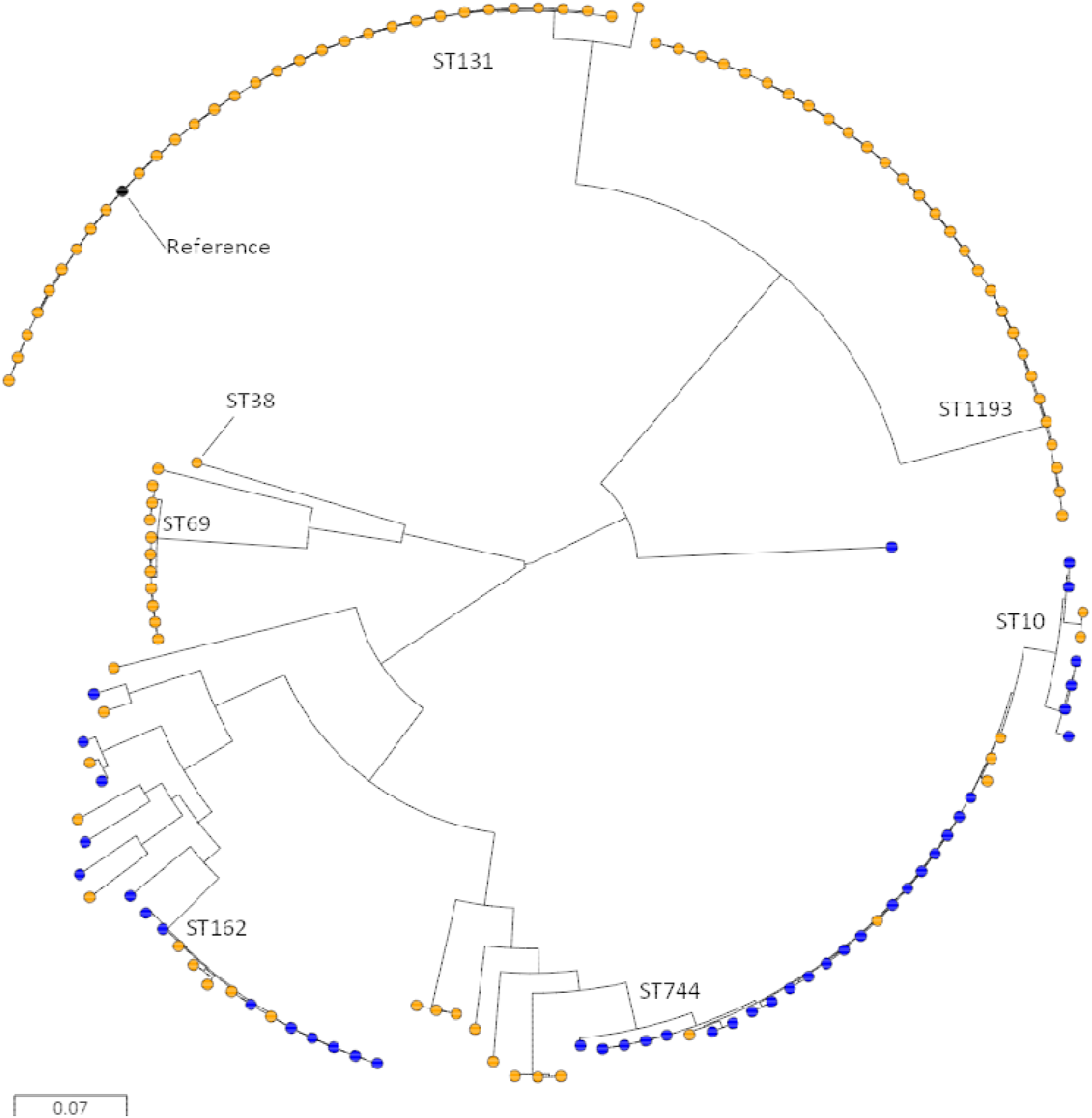
Maximum likelihood phylogenetic tree showing FQ-R, 3GC-S urinary isolates (orange) and cattle isolates (blue). Sequences were aligned against an ST131 reference (black node). Prevalent STs have been indicated.

**Figure 3.**
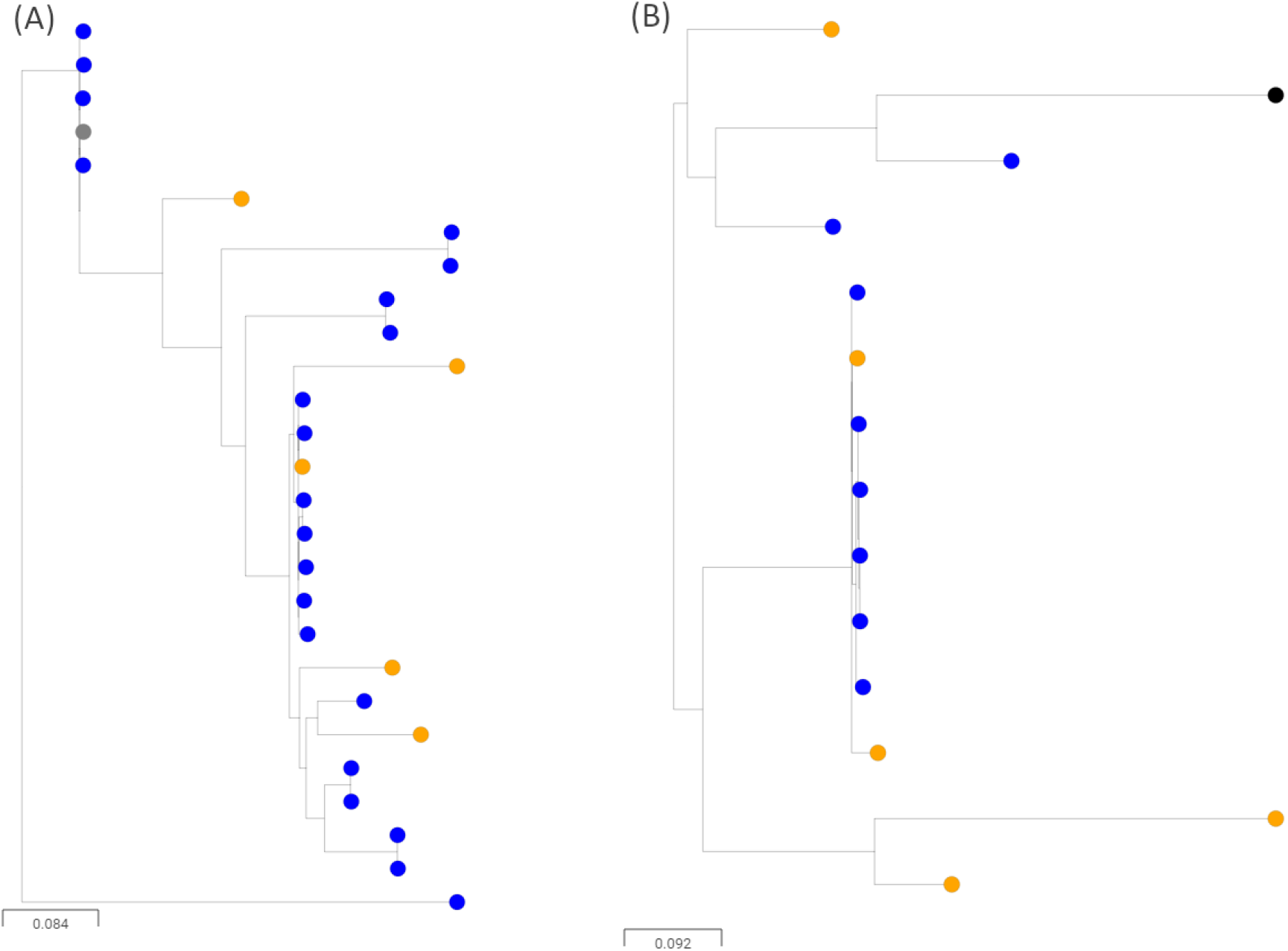
Maximum likelihood phylogenetic tree showing (A) ST744 isolates and (B) ST162 isolates also present in Figure 2. 3GC-S urinary isolates (orange) and isolates collected from environments nearby to cattle housing (blue). Sequences were aligned against references of the same ST (black node).

### Fluoroquinolone use as a driver of FQ-R E. coli on dairy farms and the suppressive effect of dry cow therapy

We next considered factors that might influence the prevalence of FQ-R *E. coli* on farms. Our aim was to identify potential interventions that might reduce FQ-R *E. coli* on farms, and so reduce any ongoing zoonotic threat. In our recent paper, we showed that the odds of a sample being positive for FQ-R *E. coli* was significantly greater if the sample came from the environment of heifer calves.^23^ This is important because dairy heifer calves are normally reared on the farm as replacement milking cows, so factors associated with the increased carriage of FQ-R *E. coli* in calves could have a long-term effect on the whole farm. Accordingly, we performed a new risk factor analysis to identify management and ABU factors associated with the odds of finding FQ-R *E. coli*-positive samples in the environment of heifer calves.

Of 631 samples collected from the environments of heifer calves, 103 (16.3%) were positive for FQ-R *E. coli*. We identified two variables that were associated with sample-level positivity for FQ-R *E. coli*. One variable - the total usage of fluoroquinolones in the year the samples were collected - was positively associated with finding FQ-R *E. coli* in a sample (odds ratio 2.39, 95% credible interval [1.01, 6.02]). Another variable - the percentage of cows within the herd dried off using any antibacterial dry cow therapy (a preparation inserted into the teats of pregnant cows thought to be at risk of developing mastitis between lactations, or with existing subclinical infections) - was negatively associated with finding FQ-R *E. coli* in samples (odds ratio 0.24, 95% credible interval [0.11, 0.50]). Full model outputs and model checking is presented in Supplementary.

## Discussion

We have shown in previous similarly powered work performed in parallel on the same study farms that there was no evidence of recent sharing of 3GC-R *E. coli* between farms and humans in the study region.^21, 22^ This agrees with the findings of similarly motivated studies from other groups.^34-36^ However, when considering FQ-R *E. coli* populations, we now report evidence for mixing of farm and human *E. coli*, and that the shared *E. coli* clones can cause bacteriuria.

A core genome SNP distance of 30 or fewer is commonly seen in phylogenetic analyses of Enterobacteriales isolates that are confirmed to be part of an acute outbreak of foodborne illness ^37^ and hospital studies frequently set a cut-off of <100 SNPs to define an outbreak.^38^ Finding FQ-R human/cattle isolates pairs differing by 71 (ST744) or 63 (ST162) SNPs is therefore suggestive of a situation where human and cattle isolates in this region do intermingle. However, this observation should be considered in the context of our finding that the closest isolates from two different farms were only three and seven SNPs apart for ST162 and ST744, respectively (**Figure 3**).

We conclude therefore, that whilst farm-related strains made up a small component of the total FQ-R urinary *E. coli* in our survey, there is evidence for zoonotic transmission to humans (**Figure 3**), which contrasts with studies considering 3GC-R *E. coli*.^21, 22, 34-36^ More work is needed to establish the exact routes of transmission, but even the minor zoonotic potential of cattle-associated FQ-R *E. coli* identified here should act as a stimulus to reduce the prevalence of such bacteria on farms. This work also suggests potential ways to achieve that objective.

We have reported at a regional level that reducing fluoroquinolone use in primary care was associated with a reduction in the proportion of *E. coli*-positive community urine samples where the isolate was FQ-R.^15^ Accordingly, it was interesting to find that overall fluoroquinolone use at farm level was positively associated with the odds of finding FQ-R *E. coli*-positive faecal samples in the environments around dairy heifer calves. The implication is that reducing fluoroquinolone use on farms may well reduce the prevalence of FQ-R *E. coli* in heifer calves, which, as they join the milking herd, may lead to a general reduction of FQ-R *E. coli* on the farm.

Notably, in mid-2018, as our surveillance of resistant *E. coli* on study farms was ending, the use of fluoroquinolones was effectively stopped, except in the very rare instance where susceptibility testing confirmed that no other antibacterial treatment option was available.^19^ This Red Tractor farm assurance scheme regulation is applicable to the vast majority of UK dairy farms.

Our final key finding was that dry cow therapy use may expedite the reduction of FQ-R *E. coli* in heifer calves, which was unexpected. We hypothesised that the reason for the association between increased usage of dry cow therapy on a farm and reduced prevalence of FQ-R *E. coli* in the environments of heifer calves was that relatively few FQ-R *E. coli* from these farms were cross-resistant to the antibacterial ingredients of dry cow therapies. These antibacterials can be released in the colostrum and first milk from treated cows. Since this colostrum is usually fed to calves at birth,^39-41^ it is plausible that calves receiving colostrum from treated cows are protected from colonisation by FQ-R *E. coli* due to the dose of antibacterial inadvertently received shortly after birth.

We calculated that 83% (in terms of weight of active ingredient) of the dry cow therapy anti-Gram-negative antibacterials used on study farms during the period of our project (2017-2018) were cephalosporins or cloxacillin (3.44 kg). Framycetin (neomycin B) made up the remainder (0.69 kg). Notably, of the 42 FQ-R cattle *E. coli* isolates subjected to WGS, only two (4.8%) were resistant to both cephalosporins and cloxacillin, as inferred from WGS.

Whilst genetically inferred framycetin resistance (presence of an *aph* gene) was more common among sequenced FQ-R cattle *E. coli* isolates (15/42, 36%), it was far less common among FQ-R isolates than among CTX-M β-lactamase-positive 3GC-R *E. coli* isolates from these same farms collected in parallel (109/135, 81% of isolates).^22^ Accordingly, there is evidence for suppression of FQ-R *E. coli* by dry cow therapy use, irrespective of active agent, because the FQ-R bacteria found on these farms are rarely resistant to the active agents used.

Much attention has been paid to reducing dry cow therapy use on dairy farms, as a way of reducing total ABU.^39-41^ We would not suggest a shift away from this approach, because inappropriate use might increase the selection of zoonotic threat of cattle *E. coli* that are resistant to other antibacterials. Indeed, we have already shown a positive association between cefquinome dry cow therapy use on dairy farms and *E. coli* carrying CTX-M type β-lactamases,^23^ the most common cause of 3GC-R in *E. coli* isolated from community UTIs in our region.^9^ Whilst use of cefquinome in dry cow therapy is also effectively stopped under the 2018 Red Tractor regulations,^19^ it is certainly possible that switching to the use of first-generation cephalosporins or cloxacillin dry cow therapy could maintain selection for CTX-M producers, which are resistant to these agents. Importantly, however, 50% of 3GC-R on our study farms was caused by chromosomally encoded AmpC hyper-production,^21, 22^ an enzyme inhibited by cloxacillin.^42^ We suggest, therefore, that a better alternative to cefquinome dry cow therapy might be cloxacillin, if all else is equal.

In contrast, a switch from cefquinome to framycetin-containing dry cow therapy might be less favoured because, as we have shown previously, framycetin dry cow therapy use co-selects CTX-M positive strains.^23^ Furthermore, whilst we have identified that framycetin resistance is less common in FQ-R *E. coli* than in *bla*_CTX-M_ positive *E. coli* on farms (36% versus 81%), we found framycetin resistance to be more common than cloxacillin resistance in FQ-R *E. coli* (36% versus 4.8%), supporting the use of cloxacillin over framycetin or a first-generation cephalosporin as the best choice to drive down FQ-R *E. coli* whilst less strongly selecting for 3GC-R *E. coli* (i.e. actively selecting against the proportion caused by AmpC enzymes).

Overall, our One Health approach to investigating selection and transmission of critically important ABR *E. coli* within our study region highlights the rare but not insignificant zoonotic potential for cattle-associated FQ-R *E. coli* as a cause of bacteriuria in humans. Our results also highlight that reducing fluoroquinolone use on farms, whilst carefully selecting the most appropriate dry cow therapy active ingredient to match the ecology of resistance found on a particular farm, should most effectively reduce that zoonotic potential. Our work certainly demonstrates the foresight of the recently introduced Red Tractor regulations designed to effectively eliminate use of highest-priority critically important antibacterials on dairy farms in the UK.^19^

## Acknowledgements

Genome sequencing was provided by MicrobesNG (http://www.microbesng.uk). We wish to thank all the farmers and veterinary surgeons who participated in this study and the biomedical scientists and other laboratory staff at Severn Pathology, Southmead Hospital, Bristol, for assistance in collecting urinary isolates. We are grateful to Dr Rob Arbon, Jean Golding Institute, University of Bristol, for advice about the Bayesian modelling work.

## Transparency declaration

D.C.B. was president of the British Cattle Veterinary Association 2018-19. Otherwise, the authors declare no competing interests. Farming and veterinary businesses who contributed data and permitted access for sample collection were not involved in the design of this study or in data analysis and were not involved in drafting the manuscript for publication.

## Author Contributions

Conceived the Study: D.C.B., K.K.R., M.B.A.

Collection of Data: O.M., H.S., J.F. E.F.P., K.M. P.N., supervised by K.B., A.P.M., T.A.C., K.K.R., M.B.A.

Cleaning and Analysis of Data: O.M., H.S., E.F.P., V.C.G., supervised by O.M.W., P.B.W., K.M.T., K.K.R., M.B.A.

Initial Drafting of Manuscript: O.M., H.S., M.B.A. Corrected and Approved Manuscript: All Authors.

