## Supplementary Information for "Limited Phylogenetic Overlap Between Fluoroquinolone-Resistant Escherichia coli Isolated on Dairy Farms and those Causing Bacteriuria in Humans Living in the Same Geographical Region"

**^1^School of Cellular & Molecular Medicine, University of Bristol, Bristol, UK**

**^2^Bristol Veterinary School, University of Bristol, Bristol, UK**

**^3^Department of Microbiology, Infection Sciences, Southmead Hospital, North Bristol NHS Trust, Bristol, UK**

**^4^Bristol Royal Infirmary, University Hospitals Bristol and Weston NHS Foundation Trust, Bristol, UK**

**Table S1**: Full results from the Bayesian analysis with mean odds ratio and 95% credible interval upper and lower limits. Effective sample size and Rhat is also presented for model evaluation.

| **Variable** | **OR** | **Lwr_CI** | **Upr_CI** | **Eff.sample_size** | **Rhat** | **Description** |
| --- | --- | --- | --- | --- | --- | --- |
| main_ceph_3_4 | 1.01 | 0.34 | 3.00 | 1944 | 1.00 | Total usage of third and fourth generation cephalosporins |
| main_fq | 2.39 | 1.01 | 6.02 | 1522 | 1.00 | Total usage of fluoroquinolones |
| main_novobiocin | 1.51 | 0.45 | 5.09 | 1510 | 1.00 | Total usage of novobiocin |
| main_total_mg_pcu | 1.57 | 0.64 | 4.18 | 1317 | 1.01 | Total usage of all antibiotics |
| reg_f_anticocc1 | 0.97 | 0.35 | 2.59 | 3439 | 1.00 | Routine use of anticoccidials |
| reg_f_anything_waste0 | 0.84 | 0.18 | 2.11 | 2419 | 1.00 | Whether waste milk was fed to any calves |
| reg_f_anything_waste1 | 1.18 | 0.46 | 5.57 | 2253 | 1.00 | Whether waste milk was fed to any calves |
| reg_f_bought_pre | 0.93 | 0.47 | 1.52 | 2671 | 1.00 | Number of cattle bought in the 12 months before the start of the project |
| reg_f_calf_housing_older1 | 0.85 | 0.22 | 1.88 | 2288 | 1.00 | Whether calves have been kept near to older animals |
| reg_f_calf_housing_type2 | 1.00 | 0.35 | 2.96 | 2822 | 1.00 | Type of calf housing |
| reg_f_calf_housing_type3 | 0.56 | 0.07 | 1.41 | 1075 | 1.00 | Type of calf housing |
| reg_f_calving_group1 | 0.88 | 0.17 | 2.45 | 2732 | 1.00 | Whether calves were born in a group pen |
| reg_f_diarrvaccY | 0.95 | 0.31 | 2.17 | 3200 | 1.00 | Routine use of calf enteric disease vaccination |
| reg_f_equine1 | 4.26 | 0.88 | 55.67 | 466 | 1.00 | Presence of horses on the farm |
| reg_f_firstmastitisCobactan | 1.01 | 0.26 | 4.25 | 3670 | 1.01 | First line mastitis tube |
| reg_f_firstmastitisMastiplanLC | 1.13 | 0.29 | 6.49 | 3389 | 1.00 | First line mastitis tube |
| reg_f_firstmastitisOrbeninLA | 1.10 | 0.30 | 7.11 | 3722 | 1.00 | First line mastitis tube |
| reg_f_firstmastitistdDmultiject | 1.19 | 0.53 | 5.28 | 2199 | 1.00 | First line mastitis tube |
| reg_f_firstmastitisUbrolexin | 0.64 | 0.03 | 2.19 | 2478 | 1.00 | First line mastitis tube |
| reg_f_firstmastitisUbroYellow | 0.80 | 0.14 | 1.91 | 2138 | 1.01 | First line mastitis tube |
| reg_f_geographical_area2 | 0.72 | 0.10 | 1.66 | 1564 | 1.00 | Which geographical area the farm is in |
| reg_f_geographical_area3 | 0.89 | 0.20 | 2.21 | 2472 | 1.00 | Which geographical area the farm is in |
| reg_f_give_col1 | 1.03 | 0.38 | 2.94 | 3429 | 1.00 | Administration of colostrum within six hours of life |
| reg_f_halocur1 | 0.61 | 0.08 | 1.45 | 1131 | 1.00 | Whether the farm routinely uses Halofuginone as a preventitive for cryptosporidiosis |
| reg_f_heifers_waste1 | 0.96 | 0.32 | 2.52 | 3160 | 1.00 | Whether waste milk has been fed to heifers |
| reg_f_herd_size | 1.07 | 0.56 | 2.44 | 3087 | 1.00 | Number of milking cows on the farm |
| reg_f_nsaiddiarr1 | 0.80 | 0.18 | 1.83 | 2258 | 1.00 | Routine use of anti-inflammatories in cases of calf diarrhoea |
| reg_f_pattern2 | 0.99 | 0.32 | 2.95 | 3283 | 1.00 | Calving pattern |
| reg_f_pneum_vacc1 | 1.54 | 0.68 | 9.27 | 1356 | 1.00 | Whether the farm routinely vaccinates calves against respiratory disease |
| reg_f_poultry1 | 0.81 | 0.18 | 1.79 | 2094 | 1.01 | Presence of poultry on the farm |
| reg_f_rain | 1.20 | 0.94 | 1.75 | 2571 | 1.00 | Average monthly rainfall |
| reg_f_scc | 0.89 | 0.42 | 1.38 | 2054 | 1.00 | Average somatic cell count |
| reg_f_time_dam.L | 1.07 | 0.53 | 2.63 | 2604 | 1.00 | Amount of time spent with dam |
| reg_f_total_cattle | 1.12 | 0.57 | 3.03 | 2244 | 1.00 | Total number of cattle on the farm |
| reg_f_trough_clean.L | 1.06 | 0.55 | 2.45 | 2641 | 1.00 | Daily cleaning of water troughs in calf housing |
| reg_f_um_spray1 | 1.11 | 0.44 | 4.14 | 3378 | 1.00 | Routine use of umbilical treatment |
| reg_f_water1 | 0.83 | 0.22 | 1.72 | 2168 | 1.00 | Water supply to the farm |
| reg_f_wean2 | 1.19 | 0.51 | 5.25 | 2395 | 1.00 | Age of weaning |
| reg_f_wean3 | 0.88 | 0.23 | 2.02 | 2983 | 1.00 | Age of weaning |
| reg_f_whichclinrespMacrolide | 1.13 | 0.55 | 3.98 | 2561 | 1.00 | First line pneumonia treatment |
| reg_f_whichclinrespPenicillimoxycillin | 1.18 | 0.35 | 10.39 | 2821 | 1.00 | First line pneumonia treatment |
| reg_f_whichclinrespTetracycline | 1.00 | 0.35 | 2.88 | 3481 | 1.00 | First line pneumonia treatment |
| reg_f_yield | 1.01 | 0.61 | 1.77 | 3345 | 1.00 | Average herd yield |
| reg_s_cefq_dct_6m1 | 0.98 | 0.34 | 2.71 | 3574 | 1.00 | Use of cefquinome dry cow therapy in the last six months |
| reg_s_ceph_dct_6m1 | 0.99 | 0.46 | 2.05 | 3268 | 1.00 | Use of cephalonium dry cow therapy in the last six months |
| reg_s_clox_dct_6m1 | 0.96 | 0.32 | 2.58 | 3227 | 1.00 | Use of cloxacillin dry cow therapy in the last six months |
| reg_s_fram_dct_6m1 | 1.32 | 0.73 | 4.55 | 2282 | 1.00 | Use of framycetin dry cow therapy in the last six months |
| reg_s_percentdryoff | 0.24 | 0.11 | 0.50 | 1644 | 1.00 | Percentage of cows that are dried off with antibiotic tubes |
| reg_s_temp | 1.16 | 0.89 | 1.73 | 2010 | 1.00 | Average monthly temperature |

### Model checking

There were no model divergencies.

#### Convergence

##### Rhat

The Rhat (Gelman-Rubin statistic) for each variable was 1.00 for all but 4 variables (which had Rhat=1.01), so we can assume the model is well converged. Trace plots (below) also provide evidence of good convergence.

##### Figure S1: Trace plots

Trace plots for the 2 risk factors identified. Variables are defined in Table S1.

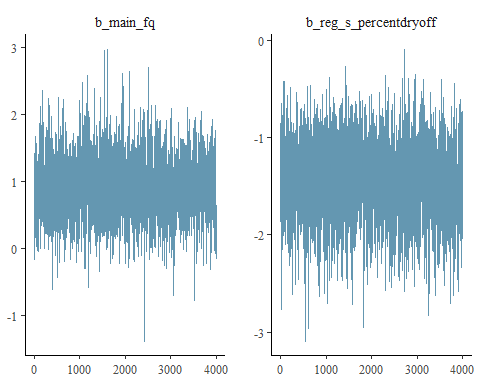

**Figure S2:** Posterior distributions

Posterior distributions for the 2 risk factors identified. Variables are defined in Table S1

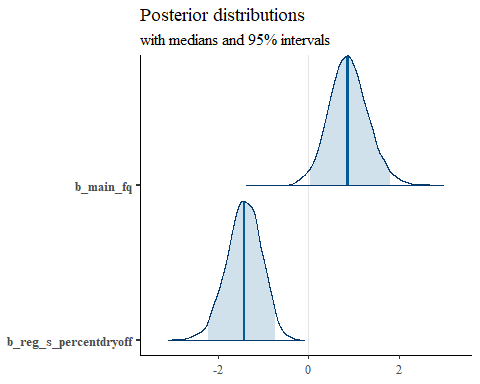

#### Table S2: Model checking.

The model was also checked using skeptical priors for the main variables whereby a mean of -0.5 and a standard deviation of 2 was applied, suggesting a narrow distribution with a negative association (which is the opposite of what was expected).

| **Variable** | **OR** | **Lwr_CI** | **Upr_CI** | **Eff.sample_size** | **Rhat** | **Description** |
| --- | --- | --- | --- | --- | --- | --- |
| main_ceph_3_4 | 0.98 | 0.34 | 2.71 | 3200 | 1.00 | Total usage of third and fourth generation cephalosporins |
| main_fq | 2.28 | 0.99 | 5.61 | 2346 | 1.00 | Total usage of fluoroquinolones |
| main_novobiocin | 1.38 | 0.43 | 4.49 | 2081 | 1.00 | Total usage of novobiocin |
| main_total_mg_pcu | 1.54 | 0.65 | 3.82 | 1925 | 1.00 | Total usage of all antibiotics |
| reg_f_anticocc1 | 0.99 | 0.34 | 2.79 | 3837 | 1.00 | Routine use of anticoccidials |
| reg_f_anything_waste0 | 0.83 | 0.16 | 2.06 | 2043 | 1.00 | Whether waste milk was fed to any calves |
| reg_f_anything_waste1 | 1.17 | 0.47 | 5.24 | 2247 | 1.00 | Whether waste milk was fed to any calves |
| reg_f_bought_pre | 0.92 | 0.45 | 1.55 | 3074 | 1.00 | Number of cattle bought in the 12 months before the start of the project |
| reg_f_calf_housing_older1 | 0.83 | 0.19 | 1.85 | 2614 | 1.01 | Whether calves have been kept near to older animals |
| reg_f_calf_housing_type2 | 0.99 | 0.33 | 2.68 | 3813 | 1.00 | Type of calf housing |
| reg_f_calf_housing_type3 | 0.59 | 0.08 | 1.44 | 1721 | 1.00 | Type of calf housing |
| reg_f_calving_group1 | 0.90 | 0.22 | 2.40 | 3172 | 1.00 | Whether calves were born in a group pen |
| reg_f_diarrvaccY | 0.96 | 0.32 | 2.40 | 4005 | 1.00 | Routine use of calf enteric disease vaccination |
| reg_f_equine1 | 4.21 | 0.86 | 50.38 | 973 | 1.01 | Presence of horses on the farm |
| reg_f_firstmastitisCobactan | 1.02 | 0.27 | 4.45 | 4837 | 1.00 | First line mastitis tube |
| reg_f_firstmastitisMastiplanLC | 1.16 | 0.32 | 8.72 | 3589 | 1.00 | First line mastitis tube |
| reg_f_firstmastitisOrbeninLA | 1.06 | 0.25 | 5.62 | 4495 | 1.00 | First line mastitis tube |
| reg_f_firstmastitistdDmultiject | 1.21 | 0.54 | 5.52 | 2577 | 1.00 | First line mastitis tube |
| reg_f_firstmastitisUbrolexin | 0.62 | 0.02 | 2.04 | 2592 | 1.00 | First line mastitis tube |
| reg_f_firstmastitisUbroYellow | 0.79 | 0.14 | 1.84 | 2548 | 1.00 | First line mastitis tube |
| reg_f_geographical_area2 | 0.71 | 0.11 | 1.61 | 1699 | 1.00 | Which geographical area the farm is in |
| reg_f_geographical_area3 | 0.89 | 0.20 | 2.22 | 2379 | 1.00 | Which geographical area the farm is in |
| reg_f_give_col1 | 1.02 | 0.40 | 2.72 | 4375 | 1.00 | Administration of colostrum within six hours of life |
| reg_f_halocur1 | 0.64 | 0.09 | 1.54 | 1971 | 1.00 | Whether the farm routinely uses Halofuginone as a preventitive for cryptosporidiosis |
| reg_f_heifers_waste1 | 0.96 | 0.32 | 2.32 | 3925 | 1.00 | Whether waste milk has been fed to heifers |
| reg_f_herd_size | 1.08 | 0.57 | 2.45 | 3380 | 1.00 | Number of milking cows on the farm |
| reg_f_nsaiddiarr1 | 0.78 | 0.17 | 1.86 | 2403 | 1.00 | Routine use of anti-inflammatories in cases of calf diarrhoea |
| reg_f_pattern2 | 0.98 | 0.31 | 2.97 | 3810 | 1.01 | Calving pattern |
| reg_f_pneum_vacc1 | 1.55 | 0.70 | 9.91 | 1754 | 1.00 | Whether the farm routinely vaccinates calves against respiratory disease |
| reg_f_poultry1 | 0.82 | 0.20 | 1.65 | 2585 | 1.00 | Presence of poultry on the farm |
| reg_f_rain | 1.20 | 0.94 | 1.74 | 2404 | 1.00 | Average monthly rainfall |
| reg_f_scc | 0.88 | 0.40 | 1.43 | 3321 | 1.00 | Average somatic cell count |
| reg_f_time_dam.L | 1.07 | 0.57 | 2.51 | 3657 | 1.00 | Amount of time spent with dam |
| reg_f_total_cattle | 1.12 | 0.62 | 2.91 | 2336 | 1.00 | Total number of cattle on the farm |
| reg_f_trough_clean.L | 1.06 | 0.54 | 2.41 | 3479 | 1.00 | Daily cleaning of water troughs in calf housing |
| reg_f_um_spray1 | 1.13 | 0.39 | 5.13 | 3259 | 1.00 | Routine use of umbilical treatment |
| reg_f_water1 | 0.82 | 0.18 | 1.82 | 2824 | 1.00 | Water supply to the farm |
| reg_f_wean2 | 1.19 | 0.51 | 5.37 | 2814 | 1.01 | Age of weaning |
| reg_f_wean3 | 0.89 | 0.24 | 2.09 | 3648 | 1.00 | Age of weaning |
| reg_f_whichclinrespMacrolide | 1.13 | 0.53 | 3.87 | 3046 | 1.00 | First line pneumonia treatment |
| reg_f_whichclinrespPenicillimoxycillin | 1.17 | 0.32 | 8.96 | 3846 | 1.00 | First line pneumonia treatment |
| reg_f_whichclinrespTetracycline | 1.00 | 0.37 | 2.54 | 3972 | 1.00 | First line pneumonia treatment |
| reg_f_yield | 1.02 | 0.62 | 1.84 | 3866 | 1.00 | Average herd yield |
| reg_s_cefq_dct_6m1 | 0.96 | 0.30 | 2.41 | 3367 | 1.00 | Use of cefquinome dry cow therapy in the last six months |
| reg_s_ceph_dct_6m1 | 0.99 | 0.43 | 2.19 | 4385 | 1.00 | Use of cephalonium dry cow therapy in the last six months |
| reg_s_clox_dct_6m1 | 0.97 | 0.34 | 2.42 | 3868 | 1.00 | Use of cloxacillin dry cow therapy in the last six months |
| reg_s_fram_dct_6m1 | 1.32 | 0.73 | 4.50 | 2422 | 1.00 | Use of framycetin dry cow therapy in the last six months |
| reg_s_percentdryoff | 0.24 | 0.11 | 0.49 | 2787 | 1.00 | Percentage of cows that are dried off with antibiotic tubes |
| reg_s_temp | 1.17 | 0.89 | 1.76 | 2775 | 1.00 | Average monthly temperature |
